# Sedentary lifestyle induces oxidative stress and atrophy in rat skeletal muscle

**DOI:** 10.1101/2024.09.27.615390

**Authors:** Irem Gungor-Orhan, Senay Akin, Scott K. Powers, Seda Olgaz-Bingol, Haydar Demirel

**Author notes:** Author of correspondence: Senay Akin.

## Abstract

Abundant evidence indicates that skeletal muscle is an important endocrine organ that plays a key role in regulating metabolic homeostasis. Therefore, maintaining healthy skeletal muscles is essential to good health. While it is established that prolonged skeletal muscle inactivity results in both oxidative stress and muscle atrophy, it remains unknown if the change from an active to a sedentary lifestyle promotes a similar increase in oxidative stress and locomotor muscle atrophy. We tested the hypothesis that the transition from active to sedentary living results in the rapid development of oxidative damage and fiber atrophy in locomotor skeletal muscles. Adult Wistar rats were randomly divided into control (CON; n=8) and sedentary (SED; n=8) groups. During a seven-day experimental period, animals in the CON group were housed in standard size cages permitting free movement, while animals in the SED group were housed in small cages that promoted sedentary behavior. At the end of the experimental protocol, soleus muscles were removed and the activities of antioxidant enzymes superoxide dismutase (SOD), catalase (CAT), and glutathione peroxidase (GPX) were determined along with two biomarkers of muscle oxidative stress (i.e., levels of advanced protein oxidation (AOPP) and 4-hydroxynonenal (4-HNE)). Living in the reduced-size cage resulted in muscle atrophy as evidenced by a 17.2% decrease in the soleus to body weight ratio (*P<0*.*001*). Moreover, the activities of SOD, CAT and GPX in the soleus muscle were significantly decreased in SED animals compared to CON (*P*<0.05). Finally, compared to CON, sedentary living increased both AOPP and 4-HNE levels in muscles of SED animals (*P*<0.001 and *P* <0.05, respectively). These findings provide the first evidence that the transition from an active to sedentary lifestyle results in the rapid onset of oxidative stress and atrophy in locomotor skeletal muscles. Importantly, these results suggest that an impaired antioxidant defense contributes to sedentary behavior-induced oxidative stress in locomotor muscles.

## Introduction

Prolonged periods of skeletal muscle disuse result in the rapid development of oxidative damage in muscle fibers leading to accelerated proteolysis and atrophy. This inactivity-induced impairment in muscle health is significant because skeletal muscle is a key endocrine organ that plays an important role in regulating metabolic homeostasis (Karstoft & Pedersen, 2016; Pedersen, 2009, 2010). Indeed, physical inactivity is linked to numerous chronic diseases including type 2 diabetes (Booth et al., 2012).

To date, most studies investigating the impact of skeletal muscle inactivity on muscle fiber size and function have employed preclinical models of muscle disuse including denervation, limb immobilization, and hindlimb unloading. Each of these models of skeletal muscle disuse confirms that prolonged muscle disuse promotes the rapid development of oxidative stress and muscle atrophy (Min et al., 2011; Muller et al., 2007; Nuoc et al., 2017; O’Leary & Hood, 2009; Powers et al., 2005). Although each of these experimental models has clinical relevance, none of these paradigms mimic the impact that physical inactivity or a sedentary lifestyle has on skeletal muscles. In this regard, it is important to appreciate that physical inactivity and sedentary lifestyle are not the same. Specifically, physical inactivity refers to individuals that do not perform the recommended amount of physical activity. In contrast, a sedentary lifestyle describes individuals that remain sedentary for more than 6 hours per day (Chuang et al., 2024; Sedentary Behaviour Research, 2012). Sedentary behavior is defined as waking behaviors that involve low energy expenditure (i.e., <1.5 METs); common sedentary behaviors include sitting, reclining, or lying (Pate et al., 2008; Sedentary Behaviour Research, 2012). Importantly, a sedentary lifestyle is also associated with an increased risk of developing several chronic diseases (Cao et al., 2022; Chuang et al., 2024; Ekelund et al., 2019; Hamilton et al., 2014; Jingjie et al., 2022). Therefore, it is important to investigate the impact of sedentary behavior on skeletal muscle health.

Recently, a novel preclinical model of sedentary behavior for rodents has emerged that permits the study of a sedentary lifestyle on skeletal muscle health (Alabi & Akomolafe, 2022; Marmonti et al., 2017). This model confines animals in small cages that permit easy access to food and water but limits physical activity to encourage sedentary behavior. Recent studies conclude that this model better simulates sedentary behavior in humans compared to previous models of extreme muscle disuse (Alabi & Akomolafe, 2022). Therefore, this new preclinical model provides an opportunity to investigate the impact of a sedentary lifestyle on oxidative damage and atrophy of skeletal muscle fibers. Thus, to determine the effect of a sedentary lifestyle on skeletal muscle, we tested the hypothesis that the transition from active to sedentary living results in the rapid development of oxidative damage and fiber atrophy in rodent locomotor skeletal muscles.

## Materials and Methods

### Animals

Sixteen male Wistar albino rats weighing 300-350 g were housed in a temperature-controlled room (22 ± 2°C) and maintained on a regular light-dark cycle (12:12 h). After arrival at the facility, animals acclimatized to the laboratory conditions for one week before the initiation of experiments to minimize the impact of the transport stress. Animals were randomly assigned into control (CON, n=8) or sedentary (SED, n=8) groups following acclimation. CON animals were housed in standard rat cages. Animals in the SED group were kept individually for seven consecutive days in reduced-volume cages. Standard maintenance rat diet and water were provided *ad libitum*. The animals’ body weight and food intake were monitored daily during the experimental period. The experimental procedures were approved by the Hacettepe University Animal Experiments Local Ethics Committee (#2022/08-07).

### Induction of sedentary life

In the current study, we employed the model originally introduced by Marmonti et al. (2017), incorporating several modifications. Briefly, small plexiglass cages with 12×12×8 cm^3^ dimensions were designed (Figure 2). These reduced-volume cages corresponded to approximately 20% of standard rat cages, allowing the animal to move around itself and providing easy access to food and water. There were grilles on the ceiling of the cage providing air circulation for the animal to breathe comfortably. Cage cleaning and bedding change were done twice a day during the whole immobilization period by transferring the animals to pre-prepared cages of the same size to prevent free movement of the animals.

**Figure 1.**
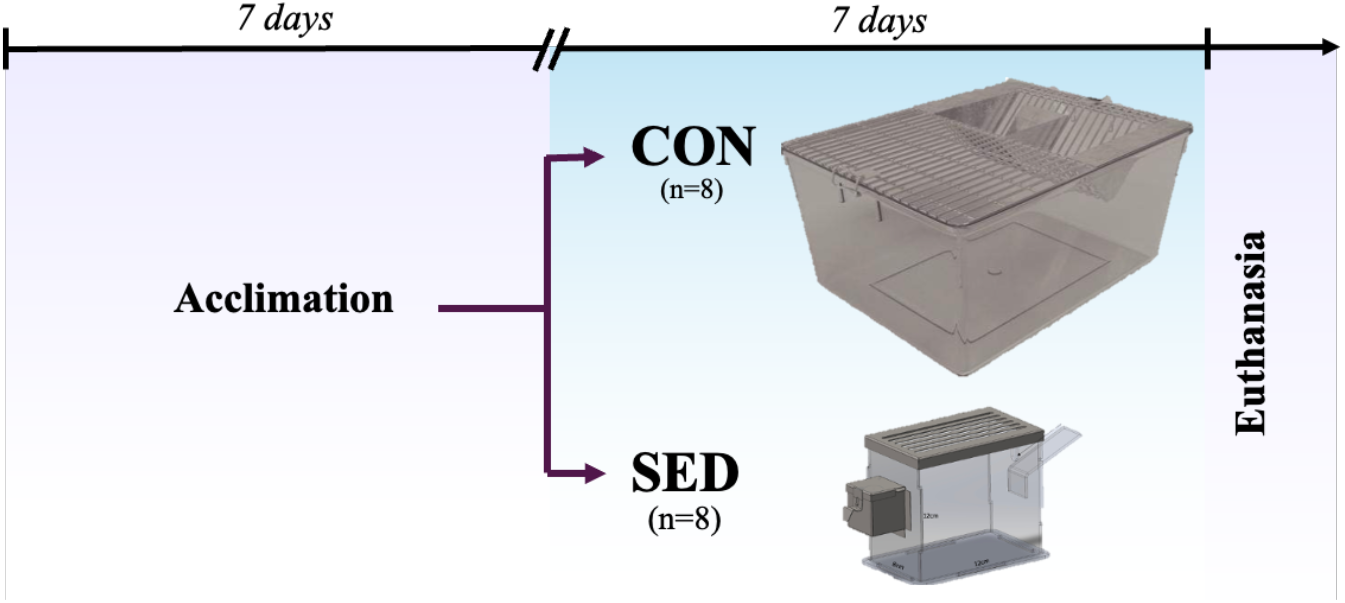
Experimental design. Following acclimation, rats were randomly assigned to CON or SED group. After seven days of the experimental protocol, rats were sacrificed on day eight.

**Figure 2.**
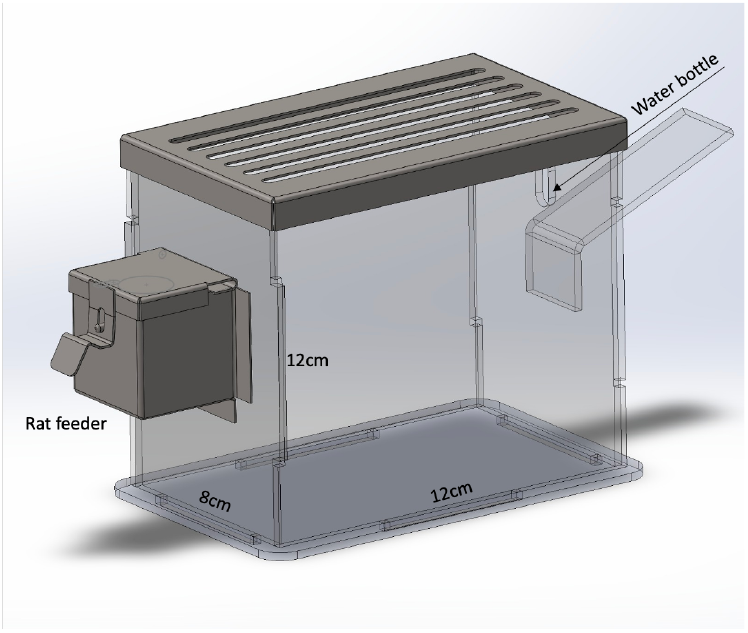
Representive picture of sedentary lifestyle model based on cage volume reduction. This model allows animals to remain immobilized, mimicking reduced physical activity conditions in humans in almost all aspects such as bed rest and sedentary lifestyle. Group SED animals were housed in small plexiglass cages with 12×12×8 cm dimensions instead of standard rat cage.

### Tissue removal and preparation

The rats were sacrificed on day eight under deep anesthesia by injecting an anesthetic cocktail solution composed of 90mg kg^-1^ ketamine and 10mg kg^-1^ xylazine. After collection of blood samples, serum samples were separated by centrifugation and aliquots were stored at −80 °C for measurement of corticosterone by liquid chromatography−tandem mass spectrometry (LC-MS/MS). Soleus muscles were trimmed from connective tissue and weighed, frozen in liquid nitrogen, and stored at -80°C until biochemical analysis.

### Determination of corticosterone levels

Serum corticosterone analyses were performed by Ultraperformance Liquid Chromatography/Mass Spectrometer (Waters Acquity Triple Quadrupole) using the Acquity UPLC BEH C18 column. Mefruside was used as an internal standard. Quantification of corticosterone is performed using selective reaction monitoring of precursors to product ion transitions, 347.2 > 329.2 m z^-1^.

### Advanced Oxidation Protein Products Content

Advanced oxidation protein products (AOPP) were determined by the spectrophotometric method as described (Witko-Sarsat et al., 1996). Briefly, thirty-milligram soleus muscle homogenized on ice in a buffer containing 5 mM Tris-HCl, 5 mM EDTA, pH, 7.4. and protease inhibitor. Homogenates were centrifuged for 10 min at 1500 *g*, 4 °C. Samples were prepared by diluting supernatants 1:8 in phosphate buffer saline. 96-well microtiter plate were loaded with 200µl of the sample, followed by 10µl 1.16M potassium iodide (KI). After incubation with KI for 2 min, the reaction was stopped by adding 20µl glacial acetic acid. The absorbance of the reaction mixture was immediately read at 340nm in a microplate reader. Chloramine T (0-100µM) standard curve was used to determine AOPP content in the samples. The concentration of AOPP was normalized to total protein content determined by bicinchoninic acid assay (BCA) assay (Takara Bio Inc.).

### Protein Isolation and Western Blotting

Fifty-milligram soleus homogenized with a buffer (5mM Tris-HCl, 5mM EDTA, pH, 7.4) containing protease inhibitors. Homogenates were centrifugated at 1500 *g* for 10 minutes at 4°C before determining protein concentration by BCA assay (Takara Bio Inc.). Levels of 4-HNE-conjugated proteins were determined by Western blot. Briefly, 33 μg of protein was loaded into sodium dodecyl sulfate polyacrylamide gel electrophoresis (SDS-PAGE). SDS-PAGE was performed on 12% gels prepared with an acrylamide:bisacrylamide ratio of 37.5:1. Proteins were separated according to their molecular weights by electrophoresis system (Bio-Rad) for approximately 2h. Then, the proteins were transferred to 0.45μm thick nitrocellulose membranes (Bio-Rad) with a semi-dry transfer system (Trans-Blot Turbo, Bio-Rad) at 25V electric current for 30 min. After the transfer process, the membranes were washed with Tris-buffered saline with 0.1% Tween 20 (TBST) and subsequently blocked with TBST containing 5% dry milk for 1 h at room temperature. Then, membranes incubated overnight with 4-HNE specific primary antibody (Abcam, #ab46545) at 4°C. After 1 h of incubation with horse radish peroxidase-conjugated secondary antibody (Cell Signaling Technology, #7074), protein bands were visualized by an enhanced chemiluminescence substrate (Biorad, #170-5061). Then, 4-HNE-conjugated protein bands between 100-25 kDa were visualized by the chemiluminescence imaging system (Chemi-Doc, Bio-Rad). Membranes were stained with Ponceau S dye to verify equal loading and transfer of proteins.

### Antioxidant Enzyme Activities

Thirty milligrams of soleus muscle was homogenized in 10 mM phosphate buffered saline, pH for CAT, GPX, and SOD activities. The homogenates were centrifuged at 10000 *g* for 10 minutes at 4°C. The supernatants were used for the determination of antioxidant enzyme activities and protein concentration of supernatants measured by BCA assay (Takara Bio Inc.) for normalization.

GPX, SOD and CAT activities were evaluated by commercial kits (Elabscience E-BC-K019-M, E-BC-K031-M, E-BC-K096-M, respectively) according to the manufacturer’s instructions. The activity of GPX was assessed through the measurement of reduced glutathione consumption. A stable yellow color, resulting from the reaction between reduced glutathione and dinitrobenzoic acid, was quantified by determining the absorbance at 412 nm. The amount of GPX activity in 1 mg of protein, catalyzing the consumption of 1 µmol L^-1^ glutathione with the subtraction of non-enzymatic reactions at 37°C for a duration of 5 minutes was defined as 1 unit. The activity of SOD was assessed through a reaction of superoxide anions (O_2_^-.^) and hydroxylamine to form nitrite, turning into purple with reaction of the chromogenic reagent. Since SOD catalyzes the conversion of superoxide anions to hydrogen peroxide (H_2_O_2_) the rection between O_2_^-.^ and hydroxylamine can be inhibited by SOD. Consequently, the SOD activity exhibited a negative correlation with the absorbance value of sample well. One unit of SOD activity is defined as the amount of SOD required to achieve a 50% inhibition ratio in a reaction solution containing 1 mg tissue protein per 1 ml. Catalase activity was assessed by the reaction of H_2_O_2_ with ammonium molybdate, resulting in the formation of a yellowish complex by measuring the optical density at 405 nm. One unit of catalase activity was defined as the quantity of catalase in 1 mg of tissue protein capable of decomposing 1 µmol of H_2_O_2_ per minute at 37°C.

### Statistical Analysis

Data for all groups are presented as the mean ± SD. All data were evaluated by the independent samples Student’s t-test. Statistical significance was set at *P* < 0.05.

## Results

This study aimed to investigate oxidative stress in skeletal muscle as a consequence of reduced physical activity, where neural stimulation and mechanical loading on skeletal muscles are reduced. The effect of a sedentary lifestyle on body weight, percentage fat mass, food, and water intake are shown in Table 1. Animals exposed to the immobilization procedure consumed similar daily food (*P* =0.055) and water (*P*=0.734) as the control group (Table 1). Seven days of sedentary living did not result in body weight differences between the experimental groups (*P* =0.063).

**Table 1.**
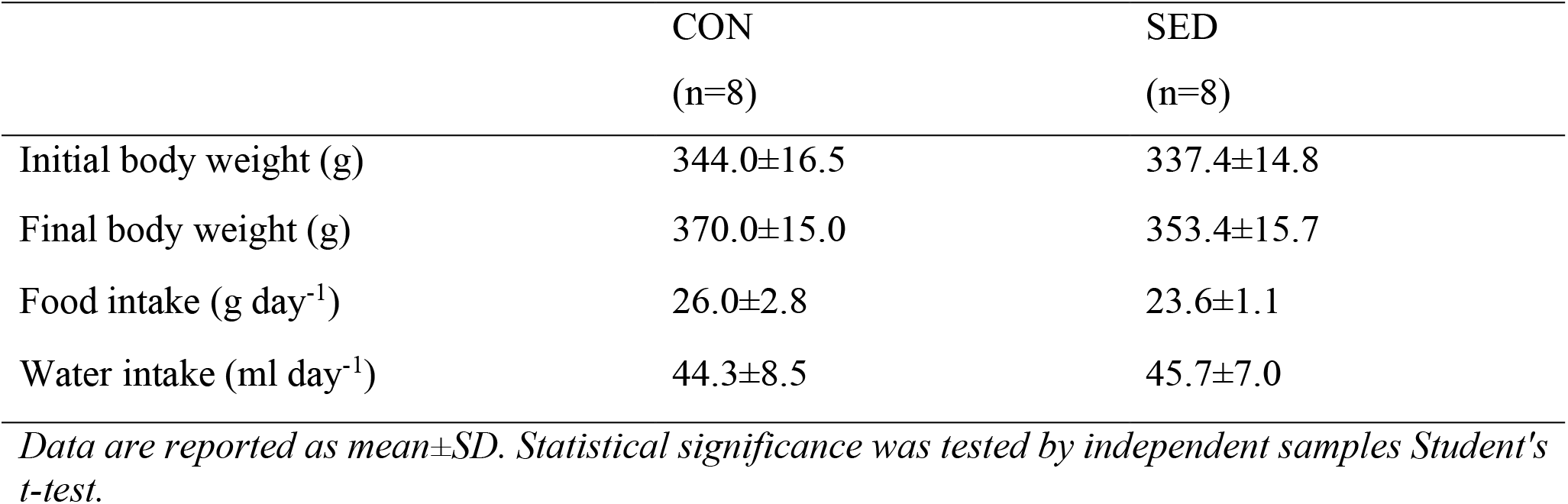
The effect of sedentary life on body weight, %fat mass, food and water intake.

|  | CON<br>(n=8) | SED<br>(n=8) |
| --- | --- | --- |
| Initial body weight (g) | 344.0±16.5 | 337.4±14.8 |
| Final body weight (g) | 370.0±15.0 | 353.4±15.7 |
| Food intake (g day<sup>-1</sup>) | 26.0±2.8 | 23.6±1.1 |
| Water intake (ml day<sup>-1</sup>) | 44.3±8.5 | 45.7±7.0 |

*Data are reported as mean±SD. Statistical significance was tested by independent samples Student's t-test.*

Table 2 shows the adrenal gland weights and serum corticosterone levels of the animals, effects of living in a reduced-volume cage for seven days in adrenal gland weight (*P*=0.574) and plasma corticosterone levels (*P*=0.057) (Table 2).

**Table 2.** Serum corticosterone levels and adrenal glands weight.

|  | CON<br>(n=8) | SED<br>(n=8) |
| --- | --- | --- |
| Corticosterone (ng ml<sup>-1</sup>) | 11.2±2.4 | 8.6±2.7 |
| Adrenal gland weight (mg 100g body weight<sup>-1</sup>) | 14.7±2.1 | 14.0±2.8 |

*Data are reported as mean±SD. Statistical significance was tested by independent samples Student's t-test.*

To determine whether immobilization causes skeletal muscle atrophy, soleus weight was measured and expressed as the soleus muscle /body weight ratio (Figure 3).

**Figure 3.**
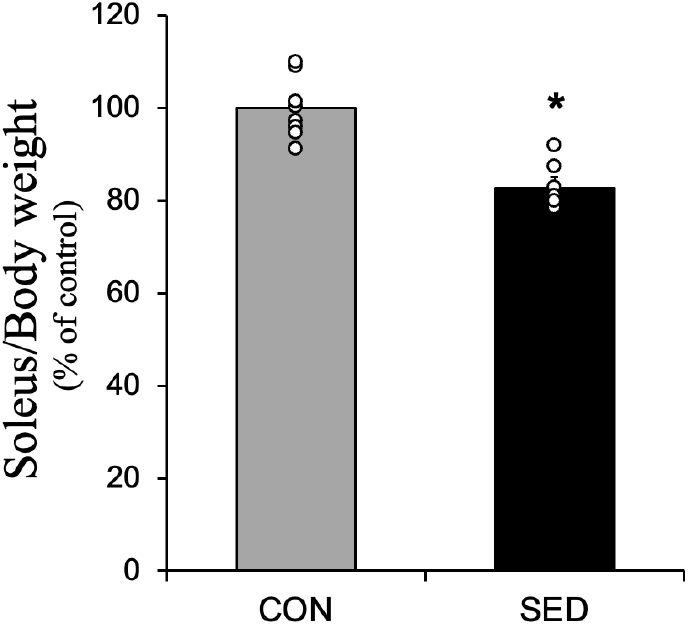
The effect of seven days of a sedentary life on soleus/body weight ratio. *Statistical difference was tested by independent samples Student’s t-test. *P* <0,001 vs. CON.

The rats housed in reduced-volume cages had 17.2% lower soleus muscle weight/body weight compared to the control group *(P* <0,001).

To investigate the effect of sedentary lifestyle on oxidative damage in soleus muscles, AOPP levels were analyzed by spectrophotometric method as an indicator of protein oxidation. The levels of the lipid peroxidation marker 4-hydroxynonenal (4-HNE) analyzed by Western Blot are shown in Figure 4.

**Figure 4.**
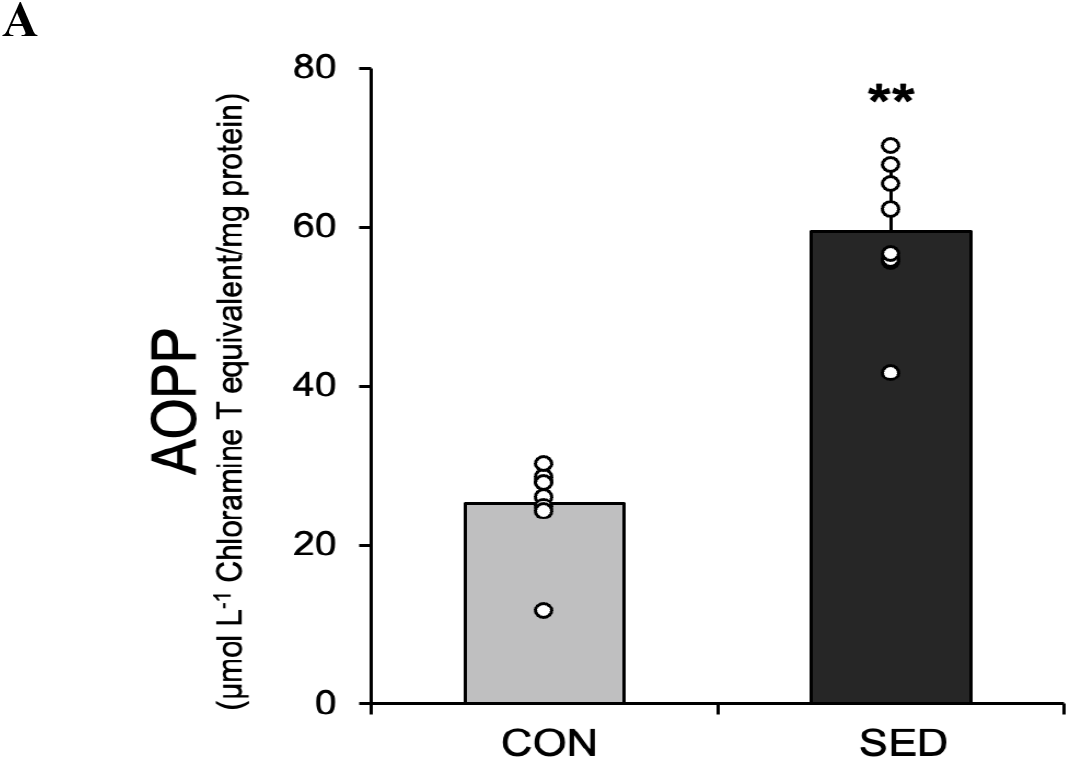
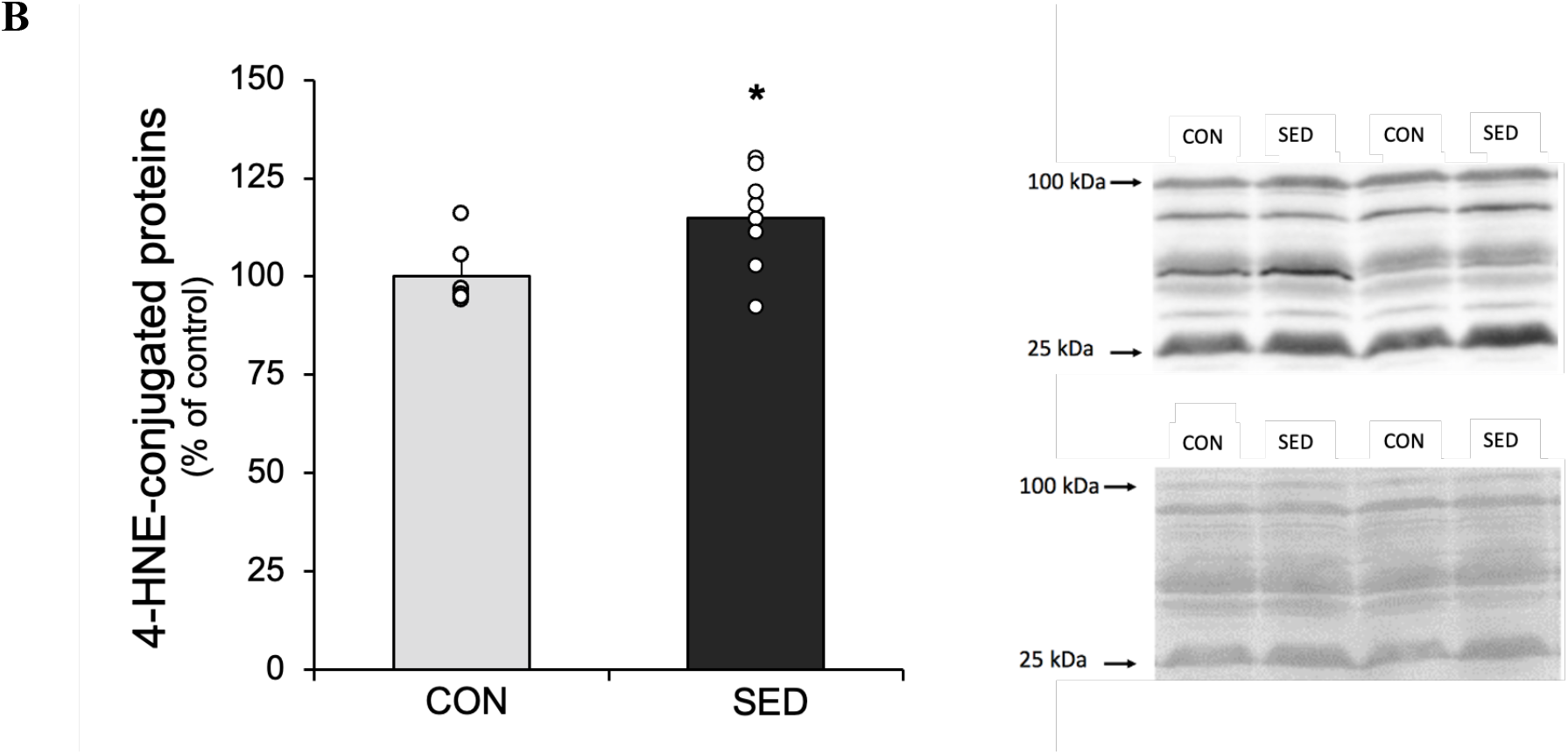
The effect of seven days of a sedentary life on oxidative damage in soleus. **A)** AOPP levels **B)** The optical density units of 4-HNE-conjugated proteins in soleus and representative Western blot for 4-HNE-conjugated proteins between 100 and 25 kDa and total protein normalization with ponceau S staining, representatively. *Statistical difference was tested by independent samples Student’s t-test*. ***\**** *P*<0,05, ***\*\*****P*<0,001 vs. CON.

Sedentary lifestyle resulted in a more than two times increase in AOPP levels in soleus muscles (Figure 4B). In addition, seven days of sedentariness increased in the 4-HNE-conjugated proteins *(P* <0,05).

To assess the impact of sedentary lifestyle on the antioxidant defense system in soleus, SOD, CAT, and GPX activities were analyzed by spectrophotometric method (Figure 5). We showed that seven days of sedentariness resulted in decreased activity of SOD, CAT, and GPX activities in the soleus (*P*<0,05) (Figure 5).

**Figure 5.**
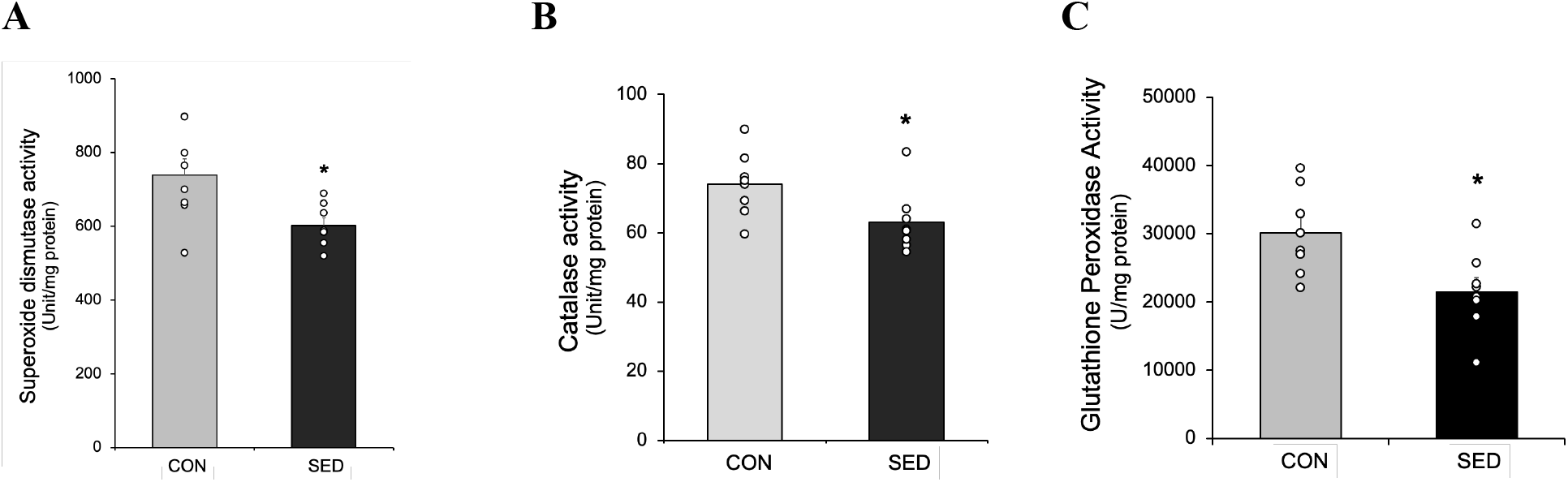
The effect of seven days of sedentary life on antioxidant enzyme activities in soleus. **A)** Superoxide dismutase activity **B)** Catalase activity **C)** Glutathione peroxidase activity. *Statistical difference was tested by independent samples Student’s t-test*. ***\**** *P*<0,05 vs. CON.

## Discussion

### Overview of major findings

To investigate the effect that a transition from active to a sedentary lifestyle has on skeletal muscle, we employed a novel preclinical model that mimics sedentary behavior in humans. Importantly, our results provide the first evidence that the transition from active to sedentary living promotes the rapid development of oxidative damage and fiber atrophy in locomotor skeletal muscles. Indeed, our findings reveal that as few as seven days of a sedentary lifestyle results in ∼17% reduction in the soleus muscle weight to body weight ratio. Notably, this sedentary living induction of muscle atrophy was associated with increased oxidative stress as evidenced by the high levels of oxidative damage biomarkers (i.e., AOPP and 4-HNE). Finally, our data also suggests that the sedentary lifestyle-induced oxidative damage in muscle may be the result of a decrease in antioxidant enzyme activity (i.e., SOD, GPX and CAT) in the muscle fibers. A brief discussion of each of these key findings follows. We begin with a critique of the preclinical experimental model.

### Critique of experimental model

Most of our knowledge about inactivity-induced locomotor muscle wasting has been garnered from rodent studies of locomotor muscle disuse using limb immobilization (e.g., casting) or hindlimb suspension. Further, studies of respiratory muscle inactivity (e.g., diaphragm) using mechanical ventilation have provided unique insight into the mechanisms responsible for inactivity-induced muscle atrophy (Falk et al., 2006; Kavazis et al., 2009; Shanely et al., 2002). Although these approaches to study muscle atrophy have clinical relevance, these models of muscle disuse and do not reflect the levels of muscle inactivity observed in humans living a sedentary lifestyle (Reidy et al., 2021)

In the current experiments using rats, we used a small cage model designed to mimic high levels of sedentary behavior in humans. This preclinical model of sedentary behavior permits the study of a sedentary lifestyle on skeletal muscle health (Alabi & Akomolafe, 2022; Marmonti et al., 2017; Reidy et al., 2021; Roemers et al., 2019; Siripoksup et al., 2024). Specifically, this model confines animals in small cages that permit free access to food and water but limits physical activity to promote sedentary behavior. A recent review concludes that, compared to previous models of extreme muscle disuse, the small cage model better simulates sedentary behavior in humans (Reidy et al., 2021); this conclusion has been supported by others (Alabi & Akomolafe, 2022; Roemers et al., 2019; Siripoksup et al., 2024).

### Sedentary lifestyle promotes skeletal muscle atrophy

Proteostasis results from a balance between the rates of protein synthesis and protein breakdown and proteostasis is essential to maintaining muscle health (Powers, Wiggs, et al., 2012). Specifically, protein abundance in muscle fibers is determined by the algebraic sum of protein synthesis minus protein breakdown. In this regard, numerous studies using a variety of models of muscle disuse conclude that inactivity-induced loss of muscle protein (i.e., atrophy) occurs due to both a decrease in muscle protein synthesis and accelerated proteolysis (Atherton et al., 2016; Bodine, 2013; Krawiec et al., 2005).

The current study demonstrates, for the first time, that the transition from an active lifestyle to a sedentary lifestyle is associated with a rapid rate of muscle atrophy. Indeed, only seven days of a sedentary lifestyle resulted in a ∼17% decrease in the soleus muscle/body weight ratio. This remarkable rate of muscle atrophy is only slightly lower than the rate of muscle atrophy observed in limb muscle disuse resulting from casting (Vazeille et al., 2008). Therefore, our findings confirm that transitioning from an active life to a sedentary behavior promotes a rapid and significant level of limb muscle atrophy.

### Transition to a sedentary lifestyle promotes rapid increases in oxidative damage to skeletal muscle fibers

Studies during the past three decades document that disuse muscle atrophy is associated with increased oxidative stress in muscle fibers (Min et al., 2011; Muller et al., 2007; Nuoc et al., 2017; O’Leary & Hood, 2009; Powers et al., 2005). Although inactivity-induced radical production likely comes several locations in the muscle fiber, evidence reveals that mitochondria are a dominant source of radical production in skeletal muscles exposed to prolonged periods of inactivity (Powers, Wiggs, et al., 2012)). Moreover, previous work in diaphragm muscle reveals that prolonged diaphragmatic inactivity during mechanical ventilation results in a diminished total antioxidant capacity (Falk et al., 2006). Similarly, the current experiments provide the first evidence that sedentary behavior results in a decrease in the activity of three key antioxidant enzymes in limb muscles. Specifically, our results reveal that only seven days of sedentary living results in significant decreases in the activities of SOD, GPX, and CAT. This observation is important because SOD is responsible for dismutation of the superoxide radical into the non-radical oxidant, hydrogen peroxide (H_2_O_2_) (Fridovich, 1997). Further, GPX and CAT play important roles in eliminating H_2_O_2_ from the cell to prevent oxidation of proteins and lipids. Together, these changes in key antioxidant enzymes reveal that sedentary living has a negative impact on the muscle antioxidant capacity.

This sedentary living-induced decrease in the total antioxidant capacity in muscle fibers can significantly impact muscle atrophy. Indeed, evidence reveals that oxidant-mediated redox signaling influences signaling pathways that regulate both protein synthesis and proteolysis in skeletal muscle fibers (reviewed in (Powers et al., 2011); (Powers et al., 2020); (Sartori et al., 2021) ;(Powers & Schrager, 2022)). In particular, inactivity-induced oxidative stress promotes a decrease in muscle protein synthesis and increases muscle proteolysis (reviewed in (Powers, Smuder, et al., 2012). The increase muscle protein degradation is mediated by oxidant-induced activation of all four major proteolytic systems including calpain, ubiquitin-proteasome system, autophagy, and caspases (Cohen et al., 2015; Egerman & Glass, 2014; Powers et al., 2016). Together, this acceleration of protein degradation pathways results in the rapid breakdown of skeletal muscle proteins resulting in muscle atrophy.

### Conclusions and future directions

To our knowledge, these are the first preclinical experiments to reveal that the transition from an active lifestyle to a sedentary lifestyle result in the rapid onset of both oxidative damage and atrophy in limb muscles. This is an important new finding that provides a warning about the negative impact that a sedentary lifestyle has on skeletal muscle health.

In regard to a sedentary lifestyle and muscle health, many unanswered questions remain. For example, can the negative effects of prolonged sedentary behavior be countered with short periods of intermittent exercise scattered throughout the day. Further, do regular bouts of exercise training result in a preconditioning of muscle fibers that protect against the atrophy and oxidative damage associated with prolonged sedentary behavior? Also, are mitochondrial-targeted antioxidants a potential counter measure to protect skeletal muscle against sedentary behavior-induced muscle atrophy and oxidative stress? Clearly, there is much more to be learned about this important topic.

*This project has supported by TUBITAK (Project number: 118S243)*

## Abbreviations

AOPP: advanced protein oxidation
BCA: bicinchoninic acid
CAT: catalase
CON: control
GPX: glutathione peroxidase
O_2_^-^: superoxide
H_2_O_2_: hydrogen peroxide
KI: potassium iodide
LC-MS/MS: liquid chromatography−tandem mass spectrometry
SDS-PAGE: sodium dodecyl sulfate polyacrylamide gel electrophoresis
SED: sedentary
SOD: superoxide dismutase
TBST: Tris-buffered saline with 0.1% Tween 20
4-HNE: 4-hydroxynonenal.

